# Emergence of Visual Center-Periphery Spatial Organization in Deep Convolutional Neural Networks

**DOI:** 10.1101/2020.02.19.956748

**Authors:** Yalda Mohsenzadeh, Caitlin Mullin, Benjamin Lahner, Aude Oliva

## Abstract

Research at the intersection of computer vision and neuroscience has revealed hierarchical correspondence between layers of deep convolutional neural networks (DCNNs) and cascade of regions along human ventral visual cortex. Recently, studies have uncovered emergence of human interpretable concepts within DCNNs layers trained to identify visual objects and scenes. Here, we asked whether an artificial neural network (with convolutional structure) trained for visual categorization would demonstrate spatial correspondences with human brain regions showing central/peripheral biases. Using representational similarity analysis, we compared activations of convolutional layers of a DCNN trained for object and scene categorization with neural representations in human brain visual regions. Results reveal a brain-like topographical organization in the layers of the DCNN, such that activations of layer-units with central-bias were associated with brain regions with foveal tendencies (e.g. fusiform gyrus), and activations of layer-units with selectivity for image backgrounds were associated with cortical regions showing peripheral preference (e.g. parahippocampal cortex). The emergence of a categorical topographical correspondence between DCNNs and brain regions suggests these models are a good approximation of the perceptual representation generated by biological neural networks.

## Introduction

Cortical regions along the ventral visual stream of the human brain (extending from occipital to temporal lobe) have been shown to preferentially activate to specific image categories^1^. For instance, while the fusiform gyrus shows specialization for faces^2^, the parahippocampal cortex (PHC) is more selective to spatial layout, places^3,4^ and large-size objects^5,6^. In characterizing the functional properties of these regions, Levy and colleagues (2001) discovered distinct topographical response patterns, such that face selective regions of the fusiform gyrus showed a strong preference for central visual field, while the building selective regions of PHC exhibited a peripheral selectivity bias to images of scene and large spaces^7^. Thus, while these regions show categorical selectivity to scenes or faces, their response patterns are strongest when their preferred category is presented in a topographically favorable location in the visual field. More specifically, the face selective voxels in the fusiform gyrus have a stronger response when faces are presented centrally; whereas scene-selective voxels show stronger activity to space features in the periphery^7–13^.

These topographical preferences raise questions regarding the origin of this functional organizing principles: does the way we look at faces and scenes in our natural visual world account for this bias? We most often fixate on faces bringing face-related information into our central, high acuity fovea to extract subtle visual features like facial expressions^14–16^. Places, on the other hand, are used for navigation, extending all around the visual field, thus we more readily perceive them with the peripheral visual information^10–13^.

Recently, a class of computational models, termed deep convolutional neural networks (DCNNs), inspired by the hierarchical architectures of ventral visual streams demonstrated striking similarities with the cascade of processing stages in the human visual system^17–25^. In particular, it has been shown that internal representations of these models are hierarchically similar to neural representations in early visual cortex^26^, mid-level (area v4), and high-level (area IT) cortical regions along ventral stream^23,27^ in primates and to functional magnetic resonance imaging (fMRI) and magnetoencephalography measurements in humans^19,22^. Recent efforts to look into the features learned by the artificial units of DCNNs have revealed the emergence of human interpretable concepts^28,29^. For example, Bau et al. (2017) showed while units in the earlier layers of the network learn patterns of edges, curves and texture, depending on the categorization task (e.g. object or scene categorization), units in the subsequent layers show selectivity to shapes and object parts or whole objects and spatial layout patterns that differentiate scenes^28^. Furthermore, they discovered that the networks trained on scene or object categorizations spontaneously learned concepts like face, people or body parts, that they never were trained on them explicitly. This work points to DCNNs as a useful model of the human visual system and motivates broader examination of the correspondence between human brain and layered-models.

These similarities motivated increasing applications of these models in hypothesis-testing of brain computations in vision^24,30–33^. In the current study, we asked whether these simplified artificial networks might show a topographical organization similar to human visual system raised from the statistics of our natural visual world. To test this hypothesis, we compared the representations of units in a deep neural network trained on both object and scene categorization^34^ (Hybrid-CNN) with representations from several category-selective areas of the visual hierarchy in the human brain. Given that this DCNN is trained on natural images representing the statistical distribution of visual features in the world, with a bias during learning similar to human visual experience (i.e. most faces are image-centered), we would expect activations of spatial selective nodes in DCNNs demonstrate category-specific topographical correspondences with human brain visual regions. Indeed, here we show that model units with central selectivity show stronger representational similarity to visual brain regions with a strong central-bias (e.g. fusiform gyrus), while model units with selectivity for image background are highly correlated with brain regions with peripheral bias (e.g. PHC).

## Results

To investigate the topographical representations of DCNNs, we probed an 8-layer deep neural network model (AlexNet^35^, Figure 1A) with high performance in categorizing scene and object images^34^. The network architecture consists of five convolutional layers followed by three fully connected layers (see Figure 1A). The model, termed Hybrid-CNN^34^, is well suited for the purposes of examining topographical representations as it is trained on both ImageNet, an object-centered dataset^36^ and Places, a scene-centered dataset^34^ making it proficient in both object and scene recognition. We fed this network a stimulus set consisting of 156 natural images of faces, animates bodies (animals and people), faces, objects and scenes. These 156 images were not used in training of the Hybrid-CNN.

**Figure 1.**
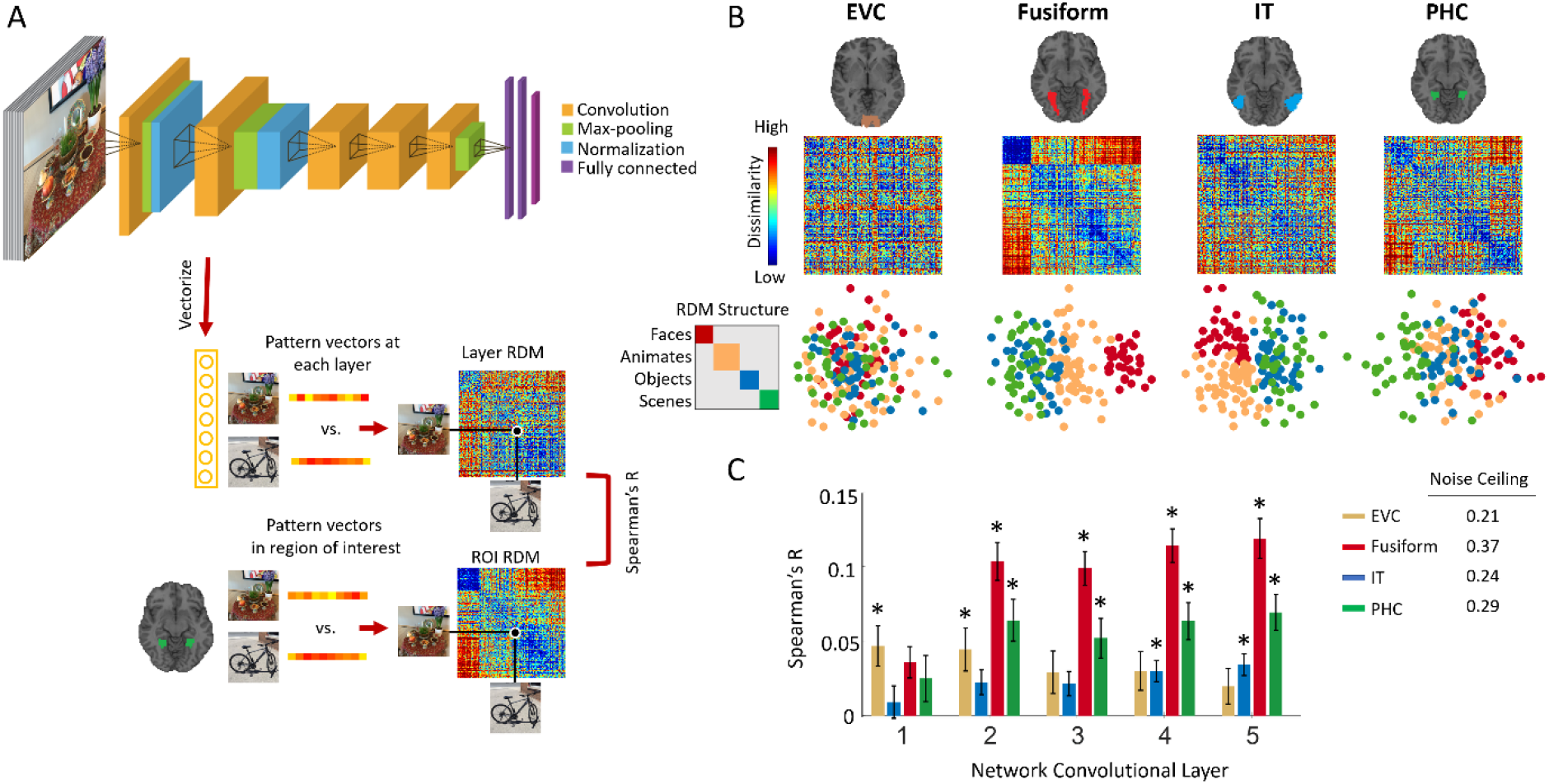
Hierarchical correspondences between layers of DCNN and brain regions of interest along ventral visual pathway. (A) For each image, the activation of units in each of the 5 convolutional layers are vectorized. RDM representation for each layer is created by computing the pairwise distance of these image specific vector patterns (1-Pearson Corr). Then fMRI RDM representations in EVC, Fusiform, IT and PHC areas are compared with the RDM representations of each convolutional layer of Hybrid-CNN by computing Spearman’s correlations. (B) Neural representations along ventral visual pathway. RDM matrices, and 2D multidimensional scaling visualization of stimuli depicted for early visual cortex (EVC), fusiform gyrus (Fusiform), inferior temporal cortex (IT) and parahippocampal cortex (PHC). (C) The correlation values for brain ROIs and layers of DCNN are depicted with bar plots. The error bars indicate the standard error of the mean and the stars above each bar indicates significant correlation above zero (N = 15, P < 0.05, Bonferroni-corrected). The noise ceiling for each brain area is reported on the right side of the panel. The pictures used in this figure are not examples of the stimulus set due to copyright.

To compare the topographical representations of the Hybrid-CNN to those from the human visual system, we collected fMRI data while participants (N = 15) viewed the 156 images in a rapid event related design and performed a vigilance task (detecting color changes in the fixation cross). The fMRI data of this study has been published in Mohsenzadeh et al. (2019)^37^.

We employed multivariate pattern analysis to resolve neural representations in functional MRI data^38–41^. To probe the topographical architecture of the visual system, we targeted four regions of interest along ventral stream which represent a range of feature and category selectivity, namely; early visual cortex (EVC), fusiform gyrus (Fusiform), inferior temporal area (IT), and parahippocampal cortex (PHC). These regions were defined anatomically according to Wang et al. (2014)^42^ and Tzourio-Mazoyer et al. (2002)^43^.

A useful tool for comparing data from different modalities has been representational similarity analysis^18,37,39,40,44–49^. This technique can quantitatively relate brain-activity measurements, such as fMRI, with computational models, such as DCNNs, by abstracting the data into representational space using matrices of pairwise similarity (a representational dissimilarity matrix-RDM). Once in the same space, we can measure the consistency of information between these two systems.

Thus, for the DCNN we abstracted the data into the representational space^18,19,22^ by extracting the layer activations, vectorized them and creating a representational dissimilarity matrix (RDM) of pairwise distances (1-Pearson corr) for each convolutional layer (1-5) as illustrated in Figure 1A. For each participant’s fMRI data, stimulus-specific voxel responses in each ROI were extracted, noise normalized^48^, arranged into pattern vectors and then the pairwise distances (1-Pearson corr) were computed and populated a 156 x 156 distance matrix, also known as representational dissimilarity matrix (RDM). In this way, the data from each source now exist in a common space.

Figure 1B depicts the subject-averaged ROI RDMs and their 2D multidimensional scaling visualizations. In line with previous literature^2–4,50^, neural representation in EVC depicts a random pattern consistent with low level visual feature processing in this area, fusiform dominantly clusters faces, IT shows a clear animate/inanimate distinction, and PHC discriminates scene images from others.

### Hierarchical correspondences between layers of Hybrid-CNN and brain regions of interest along the ventral visual pathway

Here, we first compare the hierarchical correspondences between layers of Hybrid-CNN and the ventral stream regions of interests. For this, we compute the Spearman’s rho correlations between the network layer RDMs and the individual’s ROI RDMs. Figure 1C shows comparison of fMRI representations in EVC, Fusiform, IT and PHC with the network layer-specific representations. As depicted, earlier layers of the network show significant correlation with EVC (all stats in Figure 2C, N=15; P<0.05, Bonferoni corrected). Fusiform gyrus shows progressively higher correlations along the layers of the network. IT demonstrates significant correlation with later layers (layer 4-5), consistent with a high-level object category processing in IT (for reviews see^50–52^). Finally, PHC representation is significantly correlated with mid to later layers of the network (layer 2 and 5) confirming low to high level scene semantic processes in these layers. This illustrates the distinct low to high level features of faces across the layers of this hybrid network.

**Figure 2.**
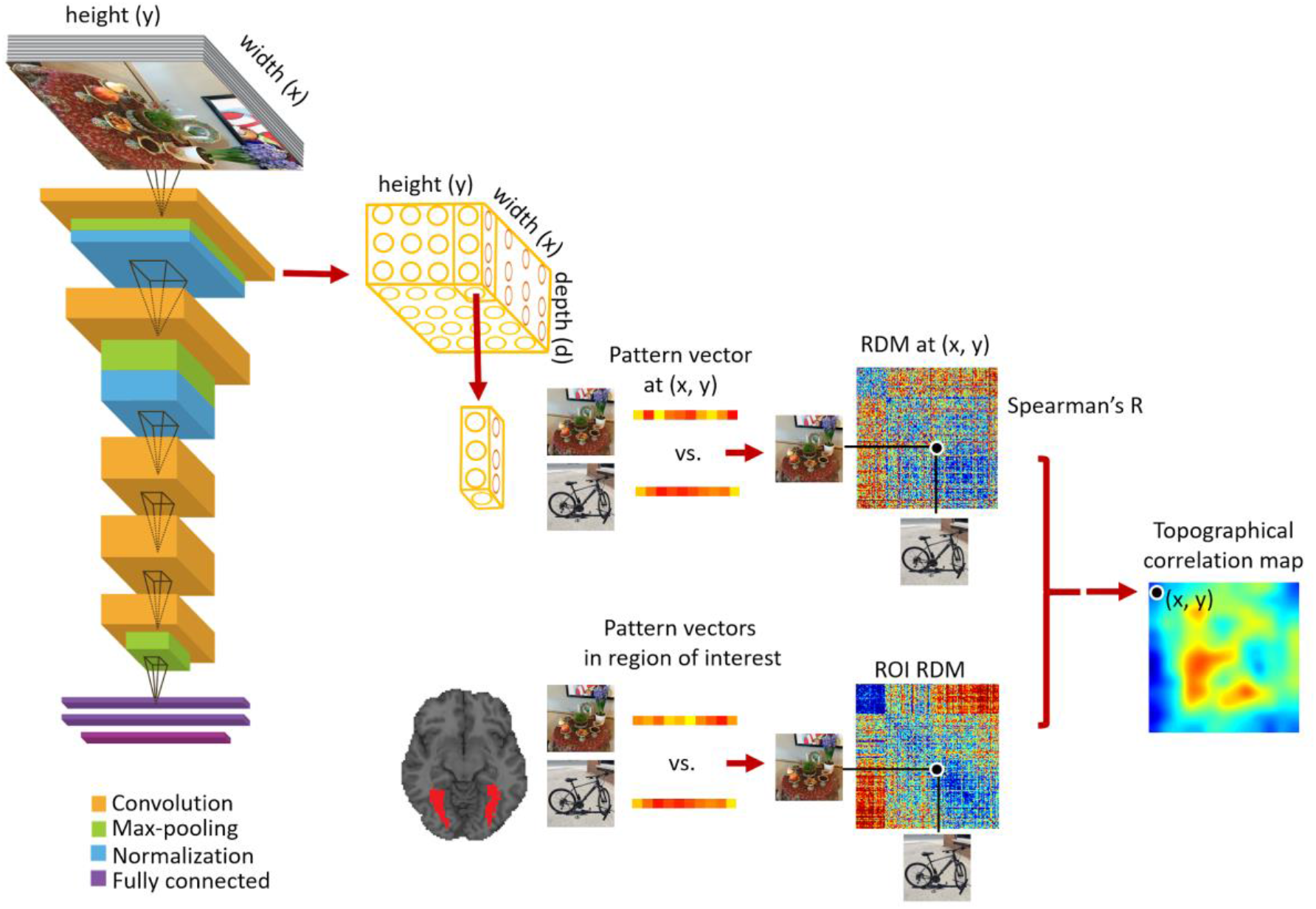
Creating topographical correlation maps. We extract the 3D activation patterns from the network convolutional layers. The first 2 Dimensions have a spatial relation with the image space (width and height). At each (x,y) position in feature maps, we extract a pattern vector with the length equivalent to the depth and construct the RDM matrix from the neural network activity patterns at each (x,y) location. Comparison of these RDM matrices with a brain ROI RDM results in a 2D correlation map which we then up-sample it to the image size (topographical map). The pictures used in this figure are not examples of the stimulus set due to copyright.

### Topographical correspondence between convolutional layers of Hybrid-CNN and human ventral visual regions

Given the hierarchical correspondence between the neural representations along ventral stream and layers of the Hybrid-CNN, we further investigated whether these representations share the same topographical central/periphery biases. We examined the spatial representations within each convolutional layer of the network and compared them with the ventral ROI representations. Figure 2 illustrates the approach: the convolutional layers of deep nets are spatially related with the 2D image space. Here, we took advantage of this correspondence and extracted the pattern vectors within each layer for pattern vectors associated with (x,y) positions in this 2D space. We used the image specific activation pattern and created RDM matrices at each (x, y) position within each layer. Then compared these RDM matrices with ROI RDMs by computing Spearman’s rho correlations resulting in 2D correlation maps as illustrated in Figure 2. We call these brain-DCNN maps *topographical correlation maps*.

Figure 3 illustrates the topographical correspondence between the five convolutional layers of the Hybrid-CNN model and the four fMRI ROIs and show the significance map corresponding to each topographical map (N=15; two-sided sign permutation tests, cluster defining threshold P<0.01, and corrected significance level P<0.05). Results are aligned with the hypothesis that the artificial network with convolutional architecture demonstrates spatial representations of center-periphery image bias highly correlated with brain regions showing similar bias: as expected, EVC, which is not category-dependent, shows a randomly distributed significant topographical pattern in the first two layers. This is consistent with previous studies showing earlier layers of network share similar representation with EVC. Furthermore, the depicted patterns indicate these low-level features are scattered over the image. Correlation maps of fusiform and layers of the network demonstrate strong center-selective patterns, consistent with the foveal-bias representations in fusiform gyrus.

**Figure 3.**
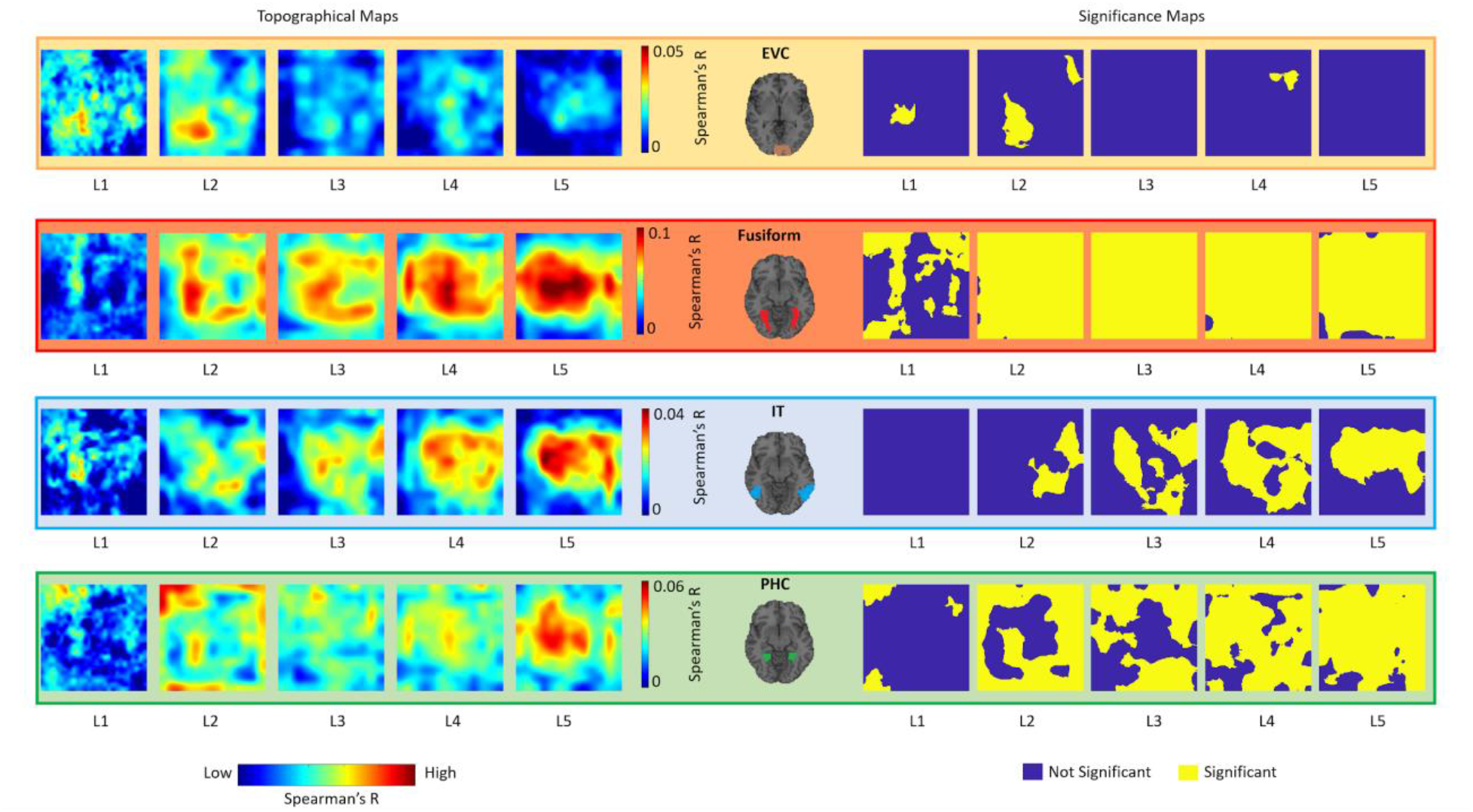
Topographical correspondence between convolutional layers of DCNNs and human ventral visual regions. For each brain-model mapping (EVC, Fusiform, IT, PHC), the first five maps show the correlational topographical maps between each convolutional layer and the brain ROI; the second five maps show the corresponding significance maps (two-sided sign permutation tests, cluster defining threshold P<0.01, and corrected significance level P<0.05). The topographical correlation maps in this figure are computed following the method depicted in Figure 2. For detailed description of RDM computations and correlations please see the Method section.

Correlation maps in IT show a dispersed pattern in mid-level layers which becomes more centralized over the layers, illustrating a mid to high-level representation transformation. Lastly, topographical maps of PHC transforms from a clear background/surrounding organization to a distributed pattern from layer 1 to 5. This suggests a transformation of low-level periphery features to scene semantics across the layers.

## Discussion

### Summary

Hierarchical correspondences have been established between primate ventral visual pathway and layers of DCNNs^19,22,23^. In the current study, using representational similarity analysis, we first replicated the previous findings showing hierarchical correspondence from low to high level ventral visual areas and layers of the deep nets (Figure 1C). Importantly, we demonstrated for the first time a topographical correspondence (central/periphery biases) between ventral brain regions and unit activations of the Hybrid-CNN (Figure 3). Our results revealed foveally biased fusiform highly correlated with unit activations of the network with the center selective visual field and peripherally biased PHC strongly correlated with unit activations of the network with periphery selective receptive fields.

### Topographical similarities of ventral visual stream and Hybrid-CNN

Visual neuroscience has focused on functional localization in establishing a set of brain areas along ventral stream that specialize in visual functions such as scene, object, face, body recognition^2–4,53–56^. On the other hand, the fields of computer vision and artificial intelligence pursue a different goal, function optimization. While artificial networks are trained on simple optimization objectives, e.g. object categorization through minimizing classification errors, they have shown emergence of similar characteristics akin to the brain areas along ventral visual pathway. Recent works have suggested that neural networks trained on natural images with categorization objectives, learn visual features along a hierarchy which matches human visual system in space and time^19,22,23,28^. For instance, deep nets trained on scene categorization showed the spontaneous emergence of object/face representations in higher layers of the network^28,29,34^. Here, we further showed that a network with convolutional constraint trained on object and scene categorization naturally demonstrates topographical correspondences with brain areas in the ventral stream showing periphery/central biases. This raises the question whether these characteristics are developed in our brain due to the position of these visual features in our visual experience (faces at the center of visual field while scenes are at the background of our vision). While neuroscientist addressed some aspects of this question^7–13^, these topographical correspondences motivate future experiments with deep neural networks to shed light on the computational principles behind these properties. Nevertheless, our findings in the current study revealing more similarities between deep nets and ventral pathway, supports two main hypotheses in vision neuroscience: first the human visual pathway optimizes the cost function of visual recognition and second the characteristics of neural tuning, internal neural representations and brain area functions along this pathway are most likely the result of this cost function optimization^57^.

### Broader implications of brain and CNN similarities

Our data reveals emergence of previously unknown similarities between the visual brain and a DCNN trained on object and scene categorization. Our results revealed a topographical correspondence between unit activations of a convolutional neural network and categorical selective brain regions. The convolutional architecture of the network limits us to investigate whether the topographical bias in the brain (and the model) is due to the statistics of the visual input or the neurons (network units) learn these characteristics. Future computational models with un-tied weights are motivated to investigate this question. We predict training such network with real world categorization tasks (objects and scenes) shapes the representation of the network units. The emergence of hierarchical as well as topographical similarities between the network and the brain implies that these characteristics of the ventral stream are most likely the result of computational objective of this pathway being visual categorization. Thus, further understanding the function, computations and connectivity architectures of visual cortex can potentially guide and give insight into brain-inspired models of vision.

## Methods

The fMRI data of this study has been published previousely^37^. Here, we briefly describe the necessary information related to experiment design and data acquisition and analysis.

### Participants

Fifteen healthy individuals (right handed; 9 females, age: mean ± SD = 27.87 ± 5.17 years) with normal or corrected to normal participated in this study. All participants signed an informed written consent form and were compensated for their time. This study was conducted in accordance with the Declaration of Helsinki and approved by the local ethics committee (Institutional Review Board of the Massachusetts Institute of Technology).

### Experiment design, task and stimulus set

Participants viewed sequence of images presented for 0.5s with 2.5s inter stimulus intervals in MRI scanner. Image trials were randomly mixed with 25% null trials during which fixation cross changed color for 100 msec and participants reported the color change by a button press. The stimulus set in our study consisted of 156 natural images of faces, animates bodies (animals and people), objects and scenes. The images were presented at the center of the screen at 6° visual angle. We acquired fMRI data in two separate sessions (each session included 5-8 runs) and images were presented once per run in random order.

### fMRI data acquisition and analysis

The fMRI data were acquired using a 3T Siemens Trio scanner with 32-channel phased-array head coil at the Athinoula A. Martinos Imaging Center at the MIT McGovern Institute for Brain Research. In each session, structural images were acquired using a standard T1-weighted sequence (176 sagittal slices, FOV = 256 mm^2^, TR = 2530 ms, TE = 2.34 ms, flip angle = 9°) and then 5-8 runs of 305 volumes of functional data were acquired for each participant (11-15 runs across the two sessions). For the functional data, gradient-echo EPI sequence was used (TR = 2000 ms, TE = 29 ms, flip angle = 90°, FOV read = 200 mm, FOV phase = 100%, bandwidth 2368 Hz/Px, gap = 20%, resolution = 3.1 mm isotropic, slices = 33, ascending interleaved acquisition).

We used SPM software to preprocess fMRI data. Functional Data of each participant were slice-time corrected, realigned and co-registered to the first session T1 structural scan, and normalized to the standard MNI space. For multivariate analysis, we used unsmoothed data. We estimated the fMRI responses to the 156 image conditions using a general linear model (GLM). We modeled the events (image conditions and nulls) with event onsets and impulse response function (duration of zero), furthermore, we included motion and run regressors in the GLM. The defined regressors were convolved with the hemodynamic response function. We then estimated the beta-values for each image condition and also the residuals from the first-level GLM. The residuals were used as an estimation of noise structure which then was employed for multivariate noise normalization (Walther et al., 2016).

In the current study, we defined four regions of interest (ROIs) along the ventral stream, early visual cortex (EVC), fusiform gyrus (Fusiform), inferior temporal cortex (IT), and parahippocampal cortex (PHC). All defined anatomically based on Wang et al. (2015)^42^ (EVC and PHC) and Tzourio-Mazoyer et al (2002)^43^ (IT and Fusiform).

### Deep convolutional neural network architecture and training

We used a deep convolutional neural network (DCNN) with architecture similar to Krizhevsky et al. (2012)^35^. This DCNN was trained both on object categories (ImageNet dataset) and scene categories (Places dataset) and called Hybrid-CNN^34^. The rationale for this choice is that our stimulus set consists of both objects and scene images and this network is trained on both ILSVRC 2012^36^ and Places dataset^34^. The network architecture includes five convolutional layers followed by three fully connected layers. For the purpose of topography comparison, we used the convolutional layers here.

### Representational similarity analysis of fMRI and DCNN units

We used representational similarity analysis to map fMRI responses and CNN units’ activations into a common space^19,22,39,41^. In this framework, the assumption is that the pairwise relationships of images are similar in the brain and the model. These pairwise relationships between 156 image-specific model/brain responses are measured in terms of dissimilarity distances (here we used 1-Pearson correlation) and summarized in a 156 × 156 representational dissimilarity matrix (RDM). We created subject-specific fMRI RDMs per region of interest (EVC, Fusiform, IT, and PHC) and model RDMs per layer or per spatial position within a convolutional layer.

In detail, in each fMRI ROI and for each of 156 image conditions we extracted the beta-value activation patterns, arranged them into vector patterns, normalized them based on the covariance of the estimated noise^48^ and then computed the pairwise dissimilarity of these 156 vector patterns by calculating 1 minus Pearson correlations. This yielded a 156×156 representational dissimilarity matrix (RDM) for each individual and ROI.

### Brain and DCNN topographical maps

In this study, we used a deep neural network by Zhou et al. (2014) called Hybrid-CNN^34^. We chose this network as it was trained both on object and scene categories and thus a suitable model to explain categories in our fMRI data. The network architecture consists of five convolutional and three fully connected layers similar to Krizhevsky et al. (2012)^35^. We extracted 3D activation maps from convolutional layers of this network for each image in our stimulus set. The first two dimensions are spatially related to the image space (width and height). As depicted in Figure 2, at each (x, y) position on the activation map, we extracted a pattern vector with the length of the depth. Then we constructed the RDM matrices from pairwise distances of these image-specific pattern vectors (1-Pearson corr). Next, we compared these neural network RDM representations with brain ROI RDMs simply by calculating the Spearman’s rho between them. This process results in a 2D correlation map for each convolutional layer which we then up-sample it to the input image dimension (topographical maps).

### Multidimensional scaling (MDS) visualization

Multidimensional scaling^58^ is an unsupervised approach to visualize the similarity relationships of conditions represented in a complex distance matrix, such that similar image conditions are visualized clustered together and different ones are depicted apart. Here, our distance matrices are 156 x 156 fMRI RDMs capturing the relations of neural patterns corresponding to 156 image stimuli in four regions of interest. The first 2 dimensions of MDS are used to visualize these images organized in 5 categories (Figure 2).

### Statistical tests

For statistical tests we used nonparametric methods with no assumption on the data distributions^59,60^. Permutation-based cluster-size inference was used for statistical inference on the topographical maps (1000 permutations, 0.01 cluster definition threshold and 0.05 cluster size threshold) with null hypothesis of zero correlation.

## Acknowledgments

Funding from the Vannevar Bush Faculty Fellowship program by the ONR (N00014-16-1-3116) and NSF award 1532591 in Neural and Cognitive Systems (to A.O.). Study conducted at the Athinoula A. Martinos Imaging Center, MIBR, MIT.

## Data availability

The data and analysis tools are available from the corresponding author upon request.

## Author contributions

Conceptualization, Y.M., A.O.; investigation, Y.M., A.O.; methodology, Y.M.; data collection, Y.M., C.M.; data analysis, Y.M., B.L.; visualization, Y.M; writing and editing, Y.M., C.M., A.O.;

## Competing Interests

The authors declare no competing interests.

